# Ancient DNA from the extinct New Zealand grayling (*Prototroctes oxyrhynchus*) reveals evidence for Miocene marine dispersal

**DOI:** 10.1101/2022.05.12.491729

**Authors:** Lachie Scarsbrook, Kieren J. Mitchell, Matthew D. McGee, Gerard P. Closs, Nicolas J. Rawlence

## Abstract

1. The evolutionary history of Southern Hemisphere graylings (Retropinnidae) in Aotearoa New Zealand—including the number of colonisation events, the directionality and timing of dispersal, and their relationship to the Australian grayling—is poorly understood. The New Zealand grayling (*Prototroctes oxyrhynchus*) is the only known freshwater fish species to have gone extinct since human arrival in New Zealand. Despite its historical abundance, only 23 formalin-fixed specimens (both wet and dried) exist in museum collections globally, which were previously non-amenable to palaeogenetic analysis.
2. Here, we used high-throughput DNA sequencing techniques, specifically designed for formalin-fixed specimens, to generate mitochondrial genomes of *P. oxyrhynchus*, and analysed these within a temporal phylogenetic framework of retropinnid and osmerid taxa.
3. We recovered strong evidence for a sister relationship between the New Zealand and Australian grayling (*P. mareana*), with the two having a common ancestor around 13.8 Mya (95% HPD: 6.1–23.2 Mya), after the height of Oligocene marine inundation in New Zealand.
4. Our temporal phylogenetic analysis suggests a single marine dispersal event between New Zealand and Australia, though the direction of dispersal is equivocal, followed by divergence into genetically and morphologically distinguishable species through isolation by distance.
5. This study provides further insights into the possible drivers of the extinction of the New Zealand grayling, and highlights how advancements in palaeogenetic techniques can be used to test evolutionary hypotheses in extinct (and living) fish, which have been comparatively neglected in the field of ancient DNA.

## 1. INTRODUCTION

The arrival of Polynesians in Aotearoa New Zealand around 1280 AD (Wilmshurst et al., 2008), followed later by Europeans (effectively the late 1700s), resulted in widespread species extinctions and habitat modification (Tennyson & Martinson, 2007; McWethy et al., 2014; Rawlence et al., 2020). Since human arrival, at least 70 species of birds have gone extinct, in addition to one mammal (Rawlence et al., 2016; Rawlence et al., 2020), one lizard (Worthy, 1987), and three frogs (Easton et al., 2021). The New Zealand grayling *Prototroctes oxyrhynchus* (or upokororo) is the only freshwater fish species known to have become extinct in New Zealand (Figure 1), with the last confirmed sighting in 1923 by the anthropologist Te Rangi Hīroa (Sir Peter Henry Buck), who captured “*over forty grayling in a funnel-shaped*” net in the Waipu River in northern New Zealand (Hīroa, 1926). Later unverified reports (e.g. newspaper articles) suggest survival into the 1950s, and species distribution models imply persistence as late as the 1970s (McDowall, 1990; Lee & Perry, 2019). However, despite their former abundance, and the intensity of 20^th^ Century fishing practices (e.g. reports of cartloads of grayling being traded; Preston, 1986), the New Zealand grayling is now only known from 23 specimens (both stuffed skins and wet-preserved) in museums in New Zealand and the United Kingdom (see McDowall, 1976). Consequently, our understanding of the New Zealand grayling’s biology, ecology, evolution, and extinction is incomplete and fragmentary.

**Figure 1.**
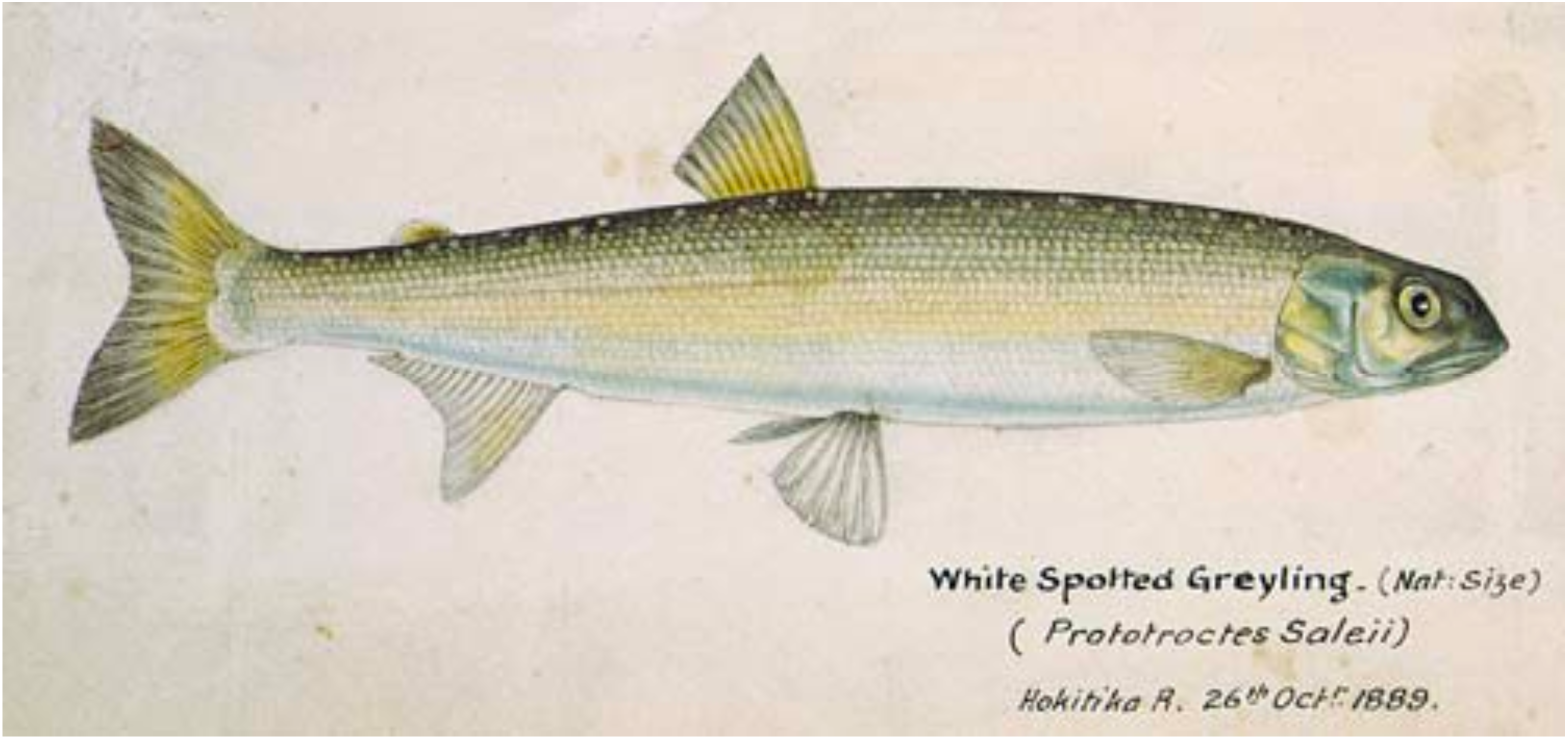
The extinct New Zealand grayling (*Prototroctes oxyrhynchus*). © Te Papa CC BY-NC-ND 4.0.

The life history traits of the New Zealand grayling are uncertain—their spawning habits are entirely unknown and they were reportedly cryptic, residing in deep pools and feeding nocturnally (McDowall, 1976). Individuals commonly reached lengths of 270– 280 mm, and weighed up to 1.36 kg (McDowall, 1990). Historically, this species inhabited both North and South Island rivers and streams proximal to the ocean (due to its amphidromous behaviour; McDowall, 1990), and may have been one of “*the most common freshwater fish in many parts of New Zealand”* (Phillips, 1923). However, despite their importance as a seasonal food source for Māori (Leach, 2006; McDowall, 2011; Fyfe & Bradshaw, 2020), skeletal remains of the New Zealand grayling have not been found in pre-European archaeological middens, though this may reflect a taphonomic artefact (McDowall, 2011; also see Seersholm et al., 2018 regarding galaxiids).

The New Zealand grayling is generally considered to be the sister-species to the endangered Australian grayling (*Prototroctes maraena*), which is presently restricted to south-eastern Australia and Tasmania. Although the genus *Prototroctes* has sometimes been attributed to a distinct family—the Prototroctidae (McDowall, 1969; McDowall, 1976; Nelson, 1994; Schwarzhans et al., 2021)—it is currently considered to belong to a subfamily (Prototroctinae) within Retropinnidae (Waters et al., 2002). The Retropinnidae (southern smelts and graylings) constitute a monophyletic group sister to the Osmeridae (northern smelts; Li et al., 2010), sharing a common ancestor ∼80 million years ago (Mya; Burridge et al., 2012). Retropinnids are endemic to southern Australia and New Zealand, and constitute three genera: *Retropinna* (southern smelts), *Prototrocte*s (southern graylings), and the monotypic New Zealand endemic *Stokellia* (Johnson & Patterson, 1996). Classification of the New Zealand and Australian graylings as congeners is based exclusively on morphological characteristics, with their division into separate species based on higher counts of lateral scale rows, vertebrae, and gill rakers in *P. oxyrhynchus* (McDowall, 1976). This hypothesis has not been tested using molecular data.

The evolutionary history of graylings in New Zealand—including the number of colonisation events, the directionality and timing of dispersal, and their relationship to Australian graylings—is poorly understood. Otoliths (calcareous structures of the inner ear) representing two species—*P. modestus* and *P. vertex*—have been recovered from the early Miocene (18.7–15.9 Mya) lacustrine deposits of the palaeolake Manuherikia, located at St Bathans (Schwarzhans et al., 2011). Identification to *Prototroctes* was based on morphological similarities to otoliths of *P. maraena*, with their relationship to *P. oxyrhynchus* unknown given the absence of known comparative material (due to the dissolution of calcareous structures in formalin and ethanol; see Supplementary Figure 1). In addition, two complete fossil fish have also been recovered from a Late Pleistocene lacustrine deposit on the north-eastern North Island (0.71–0.62 Mya), assigned to *P. oxyrhynchus* on the basis of caudal fin morphology (McDowall et al., 2006; but see Schwarzhans et al., 2011). However, as the temporal origin of *P. oxyrhynchus* is unknown, it is not possible to test whether these fossil taxa are likely to represent either stem or crown members of P*rototroctes*—the fossils from New Zealand could represent an ancient (perhaps Gondwanan) and endemic lineage that includes *P. oxyrhynchus*, or alternatively an extinct stem lineage, with *P. oxyrhynchus* descending from a recent and independent dispersal (facilitated by the juvenile marine phase of graylings).

Molecular data (i.e., ancient DNA) could help answer several questions about the taxonomic affinities and evolutionary origins of the New Zealand grayling. Unfortunately, the success of ancient DNA studies on historical ‘wet’ (i.e. preserved in fluid) and ‘dried’ (e.g. mounted) museum specimens—which encompass most teleost specimens, New Zealand grayling included—has historically been highly variable (Friedman & Desalle, 2008; Garrigos et al., 2013; Pierson et al., 2020; Pyron et al., 2022), especially due to frequent uncertainties surrounding early curatorial practices (e.g. changes to preservative fluids, most commonly unbuffered formalin to ethanol). Until recently, obtaining ancient DNA from formalin-fixed specimens has been challenging, as fixation in formalin rapidly degrades DNA (through fragmentation and base pair modification) and promotes cross-linkage both within and between DNA molecules, as well as to cellular proteins; both of which accumulate linearly with increased preservation time (Hykin et al., 2015; Hahn et al., 2021). However, methodological and technological advancements—especially the advent of high-throughput DNA sequencing—have dramatically improved success rates of obtaining genetic data from such fluid-preserved specimens. For example, Scarsbrook et al. (2022) combined DNA extraction methods optimised for ultra-short fragments (Dabney et al., 2013), single-stranded library preparation (which breaks molecular cross-links through heat-denaturation; Gansauge et al., 2017) and hybridisation capture-enrichment (González-Fortes & Paijmans, 2019) to successfully obtain complete mitochondrial genomes from historical ‘wet’ preserved New Zealand geckos.

In this study, we used paleogenetic techniques to obtain mitochondrial genome sequences from three New Zealand grayling specimens, which were compared to a newly generated mitochondrial genome from the Australian grayling, in addition to published data from other retropinnid and osmerid species. These data were used to test whether *P. oxyrhynchus* is most closely related to *P. maraena*, estimate the age of the common ancestor of *P. oxyrhynchus* and its nearest living relative, and investigate genetic structuring of *P. oxyrhynchus* populations within New Zealand.

## 2. MATERIALS AND METHODS

### 2.1 Australian grayling mitochondrial genome sequencing

Two live *P. maraena* individuals were provided via Wayne Koster (Department of Environment, Land, Water & Planning; Victoria, Australia), transported to Monash University, and euthanised with clove oil. All procedures involving live fish were performed under project AE17725 to co-author MDM at Monash University. Genomic DNA was immediately extracted from muscle tissue using a Qiagen Blood and Tissue kit, following the manufacturer’s instructions for animal tissue. DNA was sequenced by a commercial provider (Deakin Biosciences) using a PCR-free protocol for 2 × 150 bp (paired-end) sequencing on an Illumina NovaSeq S4 flow cell. The first 500,000 read pairs were extracted using bbduk.sh in the BBTools package (http:/sourceforge.net/projects/bbmap), and the mitochondrial genome assembled using tadpole.sh at a kmer size of 25. Mitochondrial genes and RNAs (rRNAs/tRNAs) were annotated using MitoFinder v. 1.4 (Allio et al., 2020) under default parameters, with the *Retropinnia semoni* (NCBI KX421785) mitochondrial genome used as a reference.

### 2.2 Australian grayling cytochrome b sequencing

Genomic DNA was extracted at the University of Otago from *P. maraena* tissue samples (fin-clips) obtained from contemporary populations in Victoria (Bunyip River; n = 5) and Tasmania (n = 1). Specifically, tissue samples (∼2 mm^2^) were added to 400 μL resuspended ‘Chelex Solution’ (2.5 g Chelex 100; 50 mL dH2O) and 2 μL Proteinase-K (20 mg/mL), and incubated at 60 °C for 24 hours at 600 rpm (on a thermoshaker). Tubes were then heated at 90 °C for 8 minutes (on a heating block), followed by centrifugation for 10 minutes at 13,000 rpm. A 1,273 bp fragment of the protein-coding gene cytochrome b (*cytb*) and associated tRNA-Thr was amplified using the following primer combination—forward: HYPSLA (5’ GTG GCT TGA AAA ACC ACC GTT; Thacker et al., 2007); reverse: Ret.Thr31 (5’ CTC CAA CCT CCG ACT TAC AAG; Page & Hughes, 2010). PCRs (10 μL) contained 1 x AmpliTaq Gold Buffer, 0.75 mM MgCl2, 0.25 mM DNTPs, 0.5 U AmpliTaq Gold DNA polymerase (ThermoFisher), 1 μM each primer (HYPSLA/Ret.Thr31) and 1 μL template DNA. Amplification was performed on an Eppendorf ProS Mastercycler (“thermocycler”) under the following conditions: an initial denaturation of 94 °C for 3 minutes; 40 cycles of 94 °C for 30 seconds, 52 °C for 1 minute and 72 °C for 2 minutes, with a final extension of 72 °C for 10 minutes. Resulting amplicons were visualised using gel electrophoresis (0.5 g agarose; 25 mL 1 x TAE Buffer; 0.5 μL SYBRsafe), and purified using ExoSap (1 μL Shrimp Alkaline Phosphatase and 1.5 μL diluted (1:20) Exonuclease I; ThermoFisher), incubated at 37°C for 30 minutes followed by 95 °C for 5 minutes on a thermocycler. Purified amplicons were bidirectionally Sanger sequenced on an ABI 3730xl DNA Analyzer (University of Otago Genetic Analysis Service) using BigDye terminator technology. Chromatogram trace files were edited (i.e., primer and poor-quality base trimming, ambiguous base-calling) in 4Peaks v. 1.8 (https://nucleobytes.com/4peaks/index.html).

### 2.3 Ancient DNA analysis of the extinct New Zealand grayling

Ancient DNA was extracted from either ‘dried’ scales or ‘wet’ tissue from historical *P. oxyrhynchus* specimens from New Zealand museum collections (n = 12; see Supplementary Table 1) using the QIAamp DNA FFPE Tissue Kit, following the manufacturer’s instructions (excluding the xylene paraffin removal step). This method utilises a heat denaturation step (at 90 °C) following tissue lysis to remove molecular cross-links, characteristic of DNA from formalin-fixed tissues (Hykin et al., 2015). Single-stranded Illumina sequencing libraries were generated from the ancient DNA extracts (following an adapted protocol; Scarsbrook et al., 2022) to enhance recovery of highly fragmented and altered DNA cross-linked to proteins (Gansauge & Meyer, 2013). Optimal cycle number (no) for the indexing of single-stranded libraries was determined through quantitative PCR (qPCR), to reduce clonality and limit heteroduplex formation (Gansauge et al., 2020). qPCRs (10 μL) were performed using 1 x Maxima® SYBR Green qPCR Master Mix (Thermo Scientific), 0.2 M each of IS7 and IS8 primers (IS7: 5’ ACA CTC TTT CCC TAC ACG AC; IS8: 5’ GTG ACT GGA GTT CAG ACG TGT) and 1 μL of diluted (1:10) library on a QuantStudio 5 Real-Time PCR System as follows: 95 °C for 10 minutes; 40 cycles of 95 °C for 30 seconds, 60 °C for 30 seconds and 72 °C for 30 seconds. Full length P5/P7 adapters containing unique 7-mer barcode combinations were added to libraries through quadruplicate indexing PCRs (to maximise complexity). Indexing PCRs (25 μL) were performed using 1 x High Fidelity PCR Buffer (Invitrogen), 2 mM MgSO4, 0.25 mM dNTPs, 1.25 U Platinum Taq DNA polymerase High Fidelity (Invitrogen), 0.5 M P5/P7 indexing primers and 5 μL library on a thermocycler as follows: 94 °C for 12 minutes; no cycles of 94 °C for 30 seconds, 60 °C for 30 seconds and 72 °C for 45 seconds; 72 °C for 10 minutes. Replicate indexed libraries were pooled, purified using AMPure XP (Agencourt) at 1.1 x bead:template ratio (following the manufacturer’s instructions) and quantified using a Qubit® dsDNA High Sensitivity Assay kit (ThermoFisher).

Endogenous DNA was selectively enriched through in-solution hybridisation-capture (following an adapted protocol; Scarsbrook et al., 2022), with biotinylated baits generated from sheared mitochondrial amplicons of the New Zealand smelt (*Retropinna retropinna*). Specifically, high-molecular weight (HMW) DNA was extracted from the high-quality tissue (liver) sample of a *R. retropinna* using the MagMAXTM DNA Multi-Sample Ultra 2.0 Kit (ThermoFisher), following the manufacturer’s instructions for saliva or whole blood. The complete *R. retropinna* mitochondrial genome (∼16.5 kb) was amplified (in a single fragment) through long-range ‘shuttle’ PCR using the following primer combination: forward: S-LA-16S-L (5’ CGA TTA AAG TCC TAC GTG ATC TGA GTT CAG; Miya & Nishida, 2000); reverse: S-LA-16S-H (5’ TGC ACC ATT RGG ATG TCC TGA TCC AAC ATC; Miya & Nishida, 2000). Long-range ‘shuttle’ PCR (50 μL) was performed using 1 x PrimeSTAR® GXL Buffer, 200 M dNTPs, 1.25 U PrimeSTAR GXL® DNA Polymerase (Takara), 0.2 M each primer (S-LA-16S-L/S-LA-16S-H) and ∼100 ng HMW DNA extract, on a thermocycler as follows: 30 cycles of 98 °C for 10 seconds and 68 °C for 16-minutes. The resulting amplicon was visualised using gel electrophoresis (0.5 g agarose; 25 mL 1 x TAE Buffer; 0.5 μL SYBRsafe) and sheared to 150-300 bp using a Picoruptor (Diagenode) sonicator with 13 cycles of: 30 seconds on, 30 seconds off, at 4 °C. Sheared amplicons were purified using AMPure XP magnetic beads (Agencourt) at 1.8 x bead:template ratio (following the manufacturer’s instructions), quantified using a Qubit® dsDNA Broad Range Assay kit (ThermoFisher), and used to generate biotinylated baits (see above).

Post-capture re-amplification PCRs (25 μL) were performed using 1 x AmpliTaq Gold Buffer (ThermoFisher), 2.5 mM MgCl2, 0.25 mM dNTPs, 1.25 U AmpliTaq Gold DNA polymerase (ThermoFisher), 0.5 M each primer (IS5: 5’ AAT GAT ACG GCG ACC ACC GA, IS6: 5’ CAA GCA GAA GAC GGC ATA CGA) and 5 μL post-capture library on a thermocycler as follows: 94 °C for 12 minutes; no cycles (as above) of 94 °C for 30 seconds, 60 °C for 30 seconds and 72 °C for 45 seconds; 72 °C for 10 minutes. Reamplified libraries were purified and quantified (as above) for equimolar pooling. Mean fragment size of the captured libraries was measured on a QIAxcel Advanced System using a 25 bp – 10 kb alignment marker; with raw data calibration and visualisation performed using QIAxcel ScreenGel Software v. 1.6.0 (Qiagen). Each library was diluted to 10 nM and run on an Illumina NextSeq (Garvan Institute of Medical Research) using 2 × 75bp (paired-end) sequencing chemistry and custom sequencing primers (CL72: 5’ ACA CTC TTT CCC TAC ACG ACG CTC TTC C; G’stein: 5’ GGA AGA GCG TCG TGT AGG GAA AGA GTG T).

Reads were demultiplexed using sabre v. 1.0 (https://github.com/najoshi/sabre) with no mismatches allowed (-m: 0). Adapter sequences were removed and paired-end reads collapsed using AdapterRemoval v. 2.3.1 (Schubert et al., 2016), with a mismatch rate of 0.33 (-mm: 3). Low-quality bases (Phred Quality Score < 30) were trimmed (--minquality: 30; --trimns) and collapsed reads shorter than 25 bp discarded (-minlength: 25). To ensure the effective removal of adapters, read quality was visualised using fastQC v. 0.11.9 (https://www.bioinformatics.babraham.ac.uk/projects/fastqc/). Mitochondrial sequence alignments were generated from collapsed reads using the BAM pipeline implemented in PALEOMIX v. 1.2.14 (Schubert et al., 2014). Briefly, collapsed reads were mapped against an Australian grayling (*Prototroctes maraena*) reference mitochondrial genome (this study) using BWA v. 0.7.17 (-n: 0.01; -o: 2; Li & Durbin, 2009). SAMtools v. 0.1.19 (Li et al., 2009) was used to select reads with a mapping Phred Quality Score >25 (-q: 25), with duplicate reads discarded in Picard v. 2.1.0 (http://broadinstitute.github.io/picard/) using the MarkDuplicates.jar tool. Misalignments from reads overlapping indels were improved using the IndelRealigner tool implemented in GATK v. 4.1.4.1 (McKenna et al., 2010). Finally, to ensure ancient DNA authenticity, characteristic damage patterns (i.e., nucleotide mis-incorporation and DNA fragmentation; Supplementary Figure 2) were assessed using MapDamage v. 2.0.8 (Ginolhac et al., 2011; Jónsson et al., 2013). Majority consensus sequences (75%) were generated for each BAM alignment in Geneious Prime, with bases only called at sites covered by ≥3 reads (with IUPAC ambiguities otherwise called). Mitochondrial genomes were annotated using the MITOS Webserver (Bernt et al., 2013), and manually verified, to determine the location and size of protein-coding genes and tRNAs/rRNAs.

### 2.4 Phylogenetic analysis

*Prototroctes oxyrhynchus* and *P. maraena* mitochondrial genomes were aligned against published retropinnid and osmerid sequences (Supplementary Table 2) using MUSCLE v. 3.8.425 (using ‘default’ parameters; Edgar, 2004) implemented in Geneious Prime v. 2021.2.2 (Biomatters; https://www.geneious.com/). Cytochrome b (1,142 bp) was extracted (in Geneious Prime) from the mitochondrial genome alignment and aligned against both modern *P. maraena* samples (n = 7; GenBank Accession Numbers: ON161129-ON161135) and published retropinnid sequences (n = 128; Supplementary Table 2). 16S rRNA (1,642 bp) was similarly extracted and aligned against published sequences (n = 15; Supplementary Table 2). Median-joining haplotype networks (Bandelt et al., 1999) were constructed from both 16S and cytb alignments in PopART (Leigh & Bryant, 2015).

Topology estimation and molecular dating were performed under a Bayesian framework in BEAST v. 1.8.4 (Drummond et al., 2012) using the mitochondrial genome alignment (subsampled to include only the most complete *P. oxyrhynchus* sequence: ON220596). Data were analysed as four partitions, using best-fitting substitution models identified using the Bayesian information criterion in PartitionFinder v. 1.0.1 (Lanfear et al., 2012): first codon positions of H-strand encoded protein coding genes (GTR+I+G), second codon positions of H-strand encoded protein coding genes (GTR+I), first codon positions of ND6 (TrN+G), and second codon positions of ND6 (K81+I). We applied a birth-death tree prior and uncorrelated lognormal relaxed clock model (with rate multiplier parameters for each partition). The continuous-time Markov chain scale reference prior (Ferreira & Suchard, 2008) was applied to the mean rate parameter and we constrained the age of the Time to Most Recent Common Ancestor (TMRCA) of the Osmeridae according to a lognormal distribution with a hard minimum at 57 Mya, mean of 13.85, and standard deviation of 1.0 (such that 95% of the prior distribution fell between 57 and 100.5 Mya)—the minimum was based on the oldest known fossil osmerid *Speirsaenigma lindoei* from the Palaeocene of Canada (Wilson & Williams, 1991), the maximum was based on the absence of crown osmerids from Late Cretaceous fossil localities, and the mean and standard deviation were chosen to reflect Burridge et al.’s (2012) posterior estimate for the equivalent node (based on a larger dataset with additional fossil age constraints). Other priors were set to their default values. We ran three Markov chain Monte Carlos (MCMCs) for 50 million generations each, sampling trees and parameters every 5,000 generations. The first 10% of each chain was removed as burn-in and remaining data were combined using LogCombiner v. 1.8.4. Tracer v. 1.7.1 (Rambaut et al., 2018) was used to monitor parameter values, ensuring convergence and effective sample sizes >200. A maximum clade credibility tree was generated in TreeAnnotator v. 1.8.4 and visualised in FigTree v. 1.4.4.

## 3. RESULTS

### 3.1 Ancient mitochondrial genomes

Mitochondrial genomes (16,591 bp) were successfully recovered from three historical New Zealand grayling (*Prototroctes oxyrhynchus*) specimens—in each instance, samples consisted of scales derived from dried specimens. Alignments produced one complete (ON220596) and two partial (ON220594–5) sequences, containing between 1,310–3,659 unique mapping reads covering 79.8–96.2% of the reference sequence to at least three-fold depth of coverage (3.6–10.8x; Supplementary Table 3). Characteristic short fragment lengths (47.7–51.4 bp; Supplementary Table 3) and damage patterns (i.e. increased C-to-T substitution frequencies at read ends) authenticated recovered DNA sequences as ancient (Supplementary Figure 2). The remaining, predominantly wet-preserved (i.e. formalin-fixed) specimens yielded very few mapped reads (0–115; Supplementary Table 3).

### 3.2 Phylogenetic analysis

Our time-calibrated Bayesian phylogeny (Fig. 2) was very well resolved, with the majority of branches receiving unequivocal support (i.e. posterior probabilities = 1.0). Divergence of the Retropinnidae and Osmeridae occurred 129.3 Mya (95% Highest Posterior Density, HPD: 83.5–183.6 Mya), with crown-ages of each family estimated at 70.4 Mya (95% HPD: 40.3–105.7 Mya) and 67.0 Mya (95% HPD: 57.2–85.9 Mya), respectively. Within Retropinnidae, the genus *Prototroctes* was the sister-taxon to *Retropinna*, with the extant *P. maraena* and extinct *P. oxyrhynchus* diverging 13.8 Mya (95% HPD: 6.1–23.2 Mya). Conversely, earlier lineage separation was inferred within the genus *Retropinna*, with divergence between *R. retropinna* and *R. semoni* estimated at 57.0 Mya (95% HPD: 30.6-87.4 Mya).

**Figure 2.**
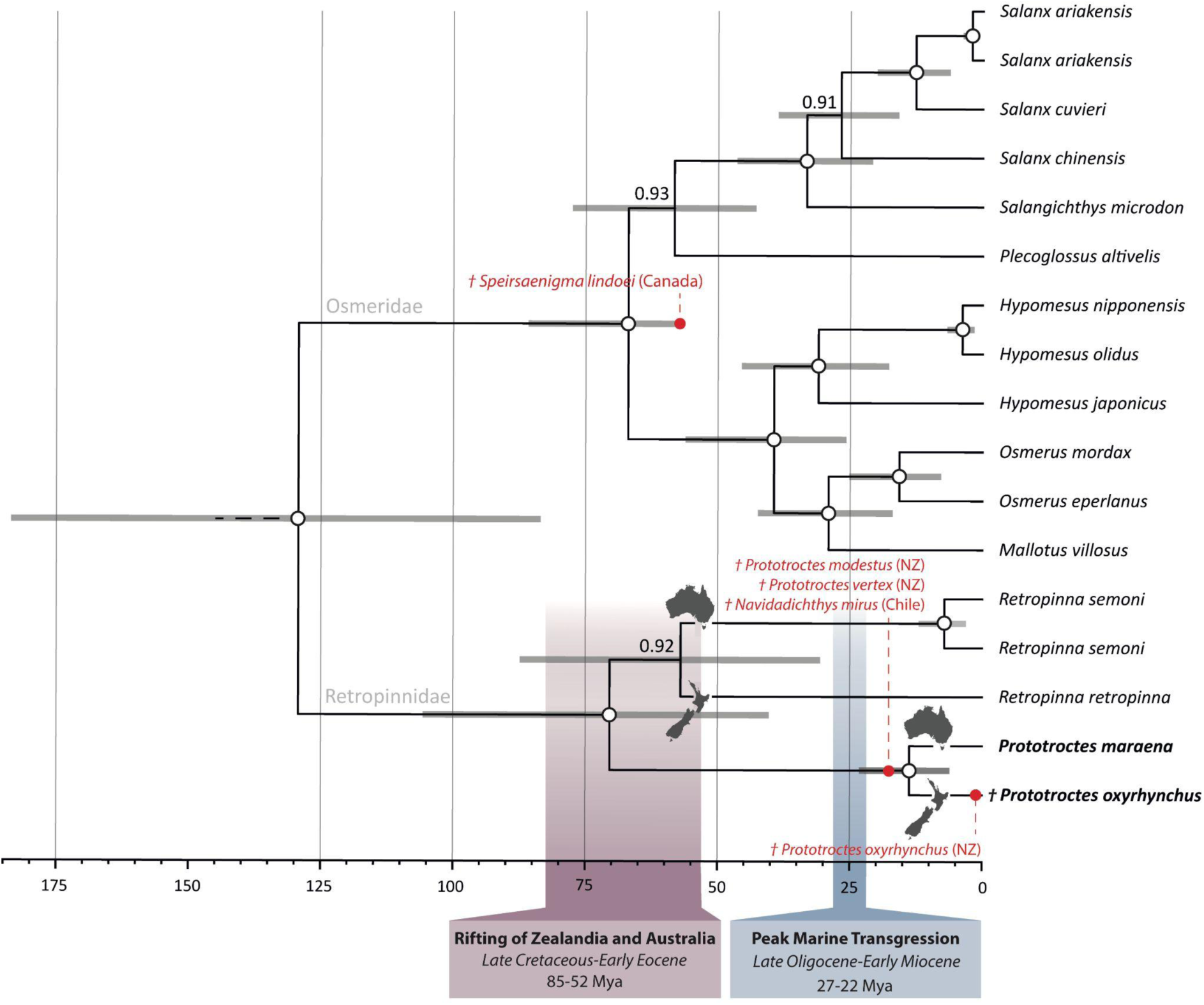
Time-calibrated Bayesian phylogeny of the Osmeriformes (Retropinnidae and Osmeridae) constructed from first and second codon positions of all 13 mitochondrial protein coding genes. 95% highest probability density (95% HPD) of the age estimate for each node is indicated by horizontal grey bars, with Bayesian posterior probabilities of 1.0 represented by white circles (otherwise values are reported). The x-axis represents time in millions of years before present (Mya). Red circles denote phylogenetic position and age of described retropinnid fossil material: *Prototroctes modestus* and *P. vertex* (18.7–15.9 Mya; Schwarzhans et al., 2011), *Navidadichthys mirus* (18-17 Mya; Schwarzhans et al., 2011), *Prototroctes oxyrhynchus* (0.71–0.62 Mya; McDowall et al., 2006)—we also indicate *Speirsaenigma lindoei* (57 Mya; Wilson & Williams, 1991), which was used to constrain the minimum age of crown Osmeridae. Timing of relevant geological events, such as rifting of the Zealandian and Australian continental blocks (Cooper & Milliner, 1993) and maximum marine inundation (i.e. ‘Oligocene drowning’; Mildenhall et al., 2014) are highlighted. Continental distribution of retropinnid lineages are indicated by outline maps. Extinct taxa are denoted †.

Cytochrome b and 16S rRNA haplotype networks (Fig. 3) revealed no evidence for population structure (or cryptic taxonomic diversity) in *P. oxyrhynchus*, with all individuals sharing the same haplotype. Conversely, two distinct haplotypes were observed in *P. maraena*, however these were not geographically structured, with both haplotypes present in the same population (for cytochrome b; Fig. 3A).

**Figure 3.**
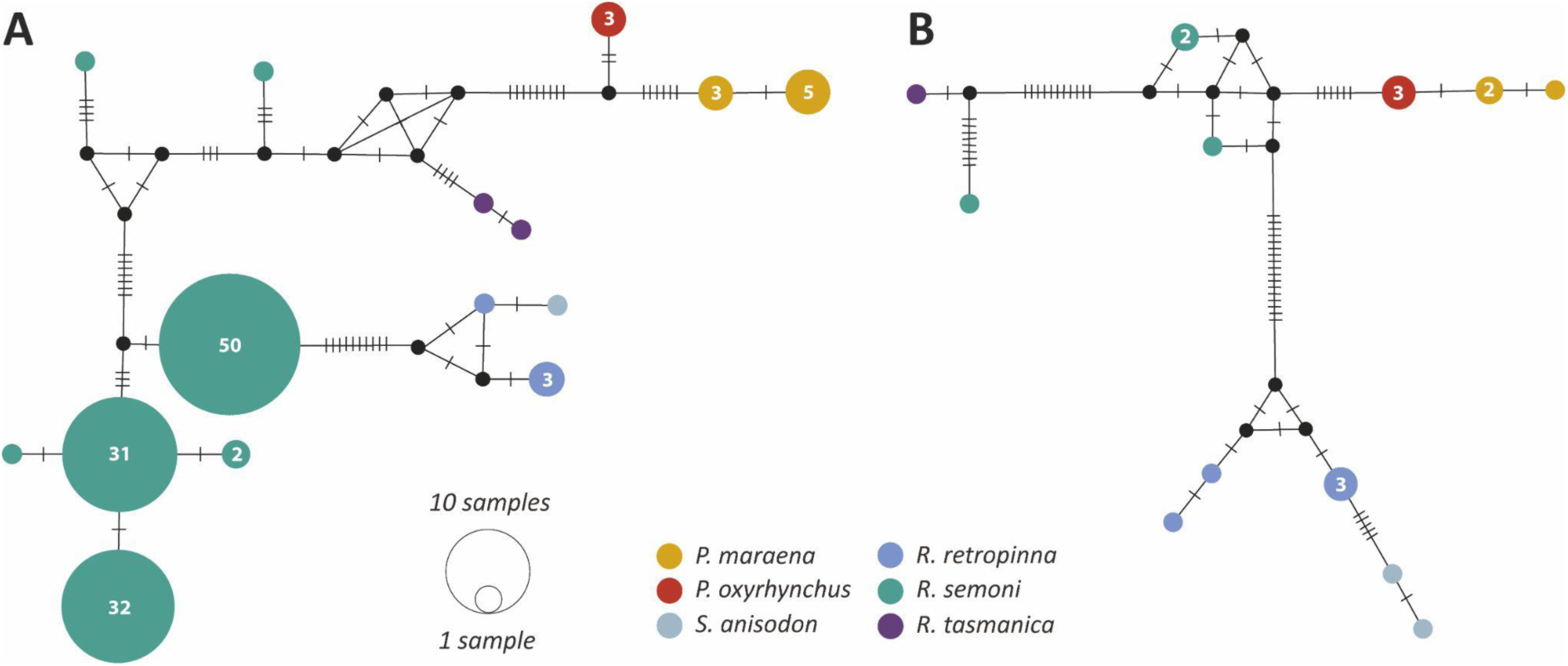
Median-joining haplotype networks of the Retropinnidae constructed from **(A)** cytochrome b (1,363 comparable sites), and **(B)** 16s rRNA (575 comparable sites) alignments in PopART (for GenBank Accession numbers, see Supplementary Table 2. Haplotypes (circles) are proportional to frequency (numbers), with number of substitutions indicated by hatches along branches. Black circles represent undetected intermediary haplotypes, with colours corresponding to species.

## 4. DISCUSSION

### 4.1 Grayling phylogeny and biogeography

Our phylogenetic results and node age estimates among extant taxa are concordant with the results of previous studies (e.g. Burridge et al., 2012; Straube et al., 2018). However, our new data allow us to include the extinct New Zealand grayling in a molecular phylogeny for the first time—our results unequivocally support a sister lineage relationship between the New Zealand and Australian graylings, and suggest that the common ancestor of the two species occurred 6.1–23.2 million years ago (Fig. 1). This divergence timing is comparable to the age of splits between osmerid sister-pairs in our analysis (see Fig. 1), and pre-dates the most-recent common ancestor of the two *R. semoni* individuals included in our alignment, which likely represent two different species (with one or both belonging to undiagnosed cryptic species; Hammer et al., 2007; Hughes et al., 2014; Schmidt et al., 2016). Consequently, it is clear that the New Zealand and Australian graylings represent two distinct species, consistent with previously observed morphological differences (McDowell, 1976). Further, extensive genetic distance between *Prototroctes* species and the remaining Retropinnidae (see Fig. 1) is consistent with previous assertions of subfamily-level (e.g. Prototroctinae)—but not family-level (e.g. Prototroctidae)—distinction of Southern Hemisphere graylings.

Extant retropinnids are restricted to south-eastern Australia and New Zealand, but a possible fossil retropinnid—with closest affinities to *Prototroctes*—has also been described from the Miocene of Chile (*Navidadichthys miru*; Schwarzhans et al., 2021). This exclusively Southern Hemisphere distribution suggests that Gondwanan vicariance may have played a role in determining the distribution of retropinnid lineages. Our estimate for the age of the common ancestor of *R. semoni* and *R. retropinna* (30.6–87.4 million years ago) is broadly consistent with this hypothesis— the distribution of these two lineages in Australia and New Zealand may have been driven by rift-related separation of these respective continental blocks through the opening of the Tasman Sea (52–85 million years ago; Cooper & Milliner, 1993). However, our results for *P. maraena* and *P. oxyrhynchus* suggest that marine dispersal, not Gondwanan vicariance, was the driver of their respective distributions in Australia and New Zealand.

While our results implicate marine dispersal in *Prototroctes*—made highly plausible by the amphidromous lifecycle of graylings—they are equivocal on the pre-Miocene distribution of the grayling lineage. Dispersing individuals could have originated from New Zealand, Australia, or even South America (since fossil data indicates this lineage was present in Chile during the Miocene). While *P. modestus* and *P. vertex* are known from New Zealand’s St Bathans fauna (Schwarzhans et al., 2011), the age of these fossil deposits (18.7–15.9 million years ago) overlaps with our estimate for the divergence between *P. maraena* and *P. oxyrhynchus*. Consequently, *P. modestus* and *P. vertex* could represent early members of the lineage leading to *P. oxyrhynchus*, and do not necessarily indicate pre-Miocene presence of graylings in New Zealand. If so, our age estimates suggest that colonisation of New Zealand by graylings might have occurred only once, after the height of the Oligocene marine transgression (27–22 million years ago) during which >80% of continental New Zealand (i.e. Zealandia) was submerged (Mildenhall et al., 2014). The extent and distribution of freshwater habitats in New Zealand would have been severely reduced (if not entirely absent) during this time, possibly resulting in the extinction and turnover of many freshwater animal lineages. Concomitant expansion of freshwater habitats in the Early Miocene (e.g. Lee et al., 2007; Schwarzhans et al., 2011) might have provided opportunities for colonisation and/or diversification that were previously limited by density-dependent founder-takes-all processes (Waters et al., 2013). Indeed, the respective common ancestors of two clades of galaxiid fishes found in New Zealand also date to the Late Oligocene or Early Miocene, where similar processes are implicated (Burridge et al., 2012; Burridge & Waters, 2020).

### 4.2 Grayling extinction

The drivers of the extinction of the New Zealand grayling are debated. Reductions in population size and geographic distribution were identified shortly after European arrival in New Zealand, with concerns raised as early as the 1870s (Rutland, 1877; Allen, 1949). By the early 1920s, the New Zealand grayling was rare and restricted to isolated rivers and streams away from human settlement (Phillips, 1923; Allen, 1949); however, it did not receive government protection until 1952, and was officially declared extinct in 2002 (Hitchmough, 2002). This decline and eventual extinction has been variously attributed to: (1) overfishing; (2) the introduction of trout, resulting in direct predation or transfer of disease (e.g. fungus, parasites), which has caused epidemics in Australian grayling populations (e.g. Johnston, 1882; Saville-Kent, 1887; Kaminskas, 2021); and/or (3) environmental modification and habitat degradation (Allen, 1949). Some authors have suggested that the disappearance of grayling from pristine unmodified rivers indicates that habitat degradation was not a primary cause of extinction (McDowall, 1990), but other authors note that source-sink dynamics could cause indirect depletion of populations even in these pristine habitats as individuals migrate to the lower density degraded habitats (Lee & Perry, 2019).

Lee and Perry (2019) suggested that up to 30% of grayling per generation could have been sustainably harvested in the absence of source-sink dynamics, reducing to only 5% when considered a single panmictic meta-population in the presence of source-sink dynamics. The presence or absence of source-sink dynamics is theoretically testable using genetic data—marked phylogeographic structure, as observed in Australian smelt (Hughes et al., 2014) and suggested by colour variation in New Zealand grayling across their range (McDowall, 1990), would imply that source-sink dynamics were not operating over large distances. We did not observe deep divergences among our three samples (Fig. 2), but these individuals were only drawn from two South Island localities (the Clutha and Hokitika Rivers)—as attempts to extract ancient DNA from other (mostly ‘wet’) grayling specimens from throughout New Zealand were not successful. However, our results suggest that additional sampling, especially of North Island individuals (e.g. Whanganui Regional Museum), combined with specialised molecular techniques for highly degraded DNA (Campos & Gilbert, 2012; Dabney et al., 2013) could be used to generate additional mitochondrial (and potentially nuclear) genomes from New Zealand grayling. These data would allow the presence of source-sink dynamics to be more effectively evaluated, and provide insight into the importance of different drivers in the extinction of the New Zealand grayling.

### 4.3 Restoring lost ecological functions to rivers by introducing Australian grayling to New Zealand

Historical reports suggest that New Zealand grayling were omnivorous, feeding on small invertebrates and algae (McDowall, 1990). As such, they likely occupied an ecological feeding niche unlike any other current extant native freshwater fish species (McDowall, 1990), and hence their (likely) extinction has resulted in some degree of lost ecological function in rivers where New Zealand grayling were formerly abundant (McDowall, 1990). Where a species translocation to restore lost ecological function is planned, the target species for translocation should be both functionally and genetically as close as possible to the species that was originally lost (Armstrong & Seddon, 2008). Based on our analysis, the Australian grayling would be the most likely extant candidate species to fulfil this criterion. That said, our work also confirms that New Zealand and Australian grayling are distinct species; hence, the translocation of Australian grayling to replace the New Zealand grayling is clearly a species introduction (i.e. ecological surrogate), not a reintroduction (Armstrong & Seddon, 2008). Whilst this point might seem to be mere semantics, the distinction does serve to highlight the potential challenges of such a management step. Given the time since their last common ancestor, the two species would have likely diverged in a variety of key morphological and behavioural traits, thus predicting how Australian grayling would respond to New Zealand conditions is impossible, both in terms of the viability of such an introduction, but also with respect to the goal of restoring lost ecological function.

## ACKNOWLEDGEMENTS

We thank Tom Trnski and Severine Hannam (Auckland War Memorial Museum); Paul Scofield (Canterbury Museum); Karen Cook (Fish and Game, Nelson); Andrew Stewart, Jeremy Barker and Clive Roberts (Museum of New Zealand Te Papa Tongarewa); Eimear Egan (National Institute of Water and Atmospheric Research); and Kane Fleury and Emma Burns (Otago Museum) for access to *P. oxyrhynchus* material for genetic analysis; James Maclaine (Natural History Museum, United Kingdom) for generating X-ray images of NMUK *P. oxyrhynchus* syntypes; Chris Burridge (University of Tasmania), Matt Jarvis (University of Otago), and Wayne Koster (Arthur Rylah Institute for Environmental Research) for access to fin-clips and otoliths from contemporary Australian grayling (*P. maraena*) populations; Jesse Wansbrough (University of Otago) for providing the *R. retropinna* tissue sample; and Alex Verry and Tania King (University of Otago) for their assistance in optimising DNA extraction, PCR, and next-generation sequencing protocols. We also acknowledge the use of New Zealand eScience Infrastructure (NeSI) high-performance computing facilities as part of this research. We acknowledge that Māori, the indigenous people of Aotearoa New Zealand, have kaitiaki (guardianship) over the organisms from their rohe (tribal area). Funding was provided by the University of Otago.

## AUTHOR CONTRIBUTIONS

Conceptualisation: GPC, NJR. Developing methods: LS, NJR. Conducting the research: LS, MDM. Data analysis: LS, KJM, MDM. Data interpretation: LS, KJM, GPC, NJR. Writing: LS, KJM, MDM, GPC, NJR.

## DATA AVAILABILITY

The data that support the findings of this study are openly available, with raw sequencing reads (.fastq files) deposited in Dryad [***]; and consensus mitochondrial genomes openly available in GenBank [Accession Numbers: ON161129-ON161135; ON220594–ON220597].

## REFERENCES

Allen, K. R. (1949). The New Zealand grayling - a vanishing species. Tuatara, 2(1), 22–26.

Allio, R., Schomaker-Bastos, A., Romiguier, J., Nabholz, B., & Delsuc, F. (2020). MitoFinder: efficient automated large-scale extraction of mitogenomic data in target enrichment phylogenomics. Molecular Ecology Resources, 20 (4), 892–905.

Armstrong, D. P., & Seddon, P. J. (2008) Direction in reintroduction biology. Trends in Ecology and Evolution, 23(1), 20–25.

Bandelt, H. J., Forster, P., & Röhl, A. (1999). Median-joining networks for inferring intraspecific phylogenies. Molecular Biology and Evolution, 16 (1), 37–48.

Bernt, M., Donath, A., Jühling, F., Externbrink, F., Florentz, C., Fritzsch, G., … & Stadler, P. F. (2013). MITOS: improved de novo metazoan mitochondrial genome annotation. Molecular Phylogenetics and Evolution, 69(2), 313–319.

Burridge, C. P., McDowall, R. M., Craw, D., Wilson, M. V., & Waters, J. M. (2012). Marine dispersal as a pre-requisite for Gondwanan vicariance among elements of the galaxiid fish fauna. Journal of Biogeography, 39(2), 306–321.

Burridge, C. P., & Waters, J. M. (2020). Does migration promote or inhibit diversification? A case study involving the dominant radiation of temperate Southern Hemisphere freshwater fishes. Evolution, 74, 1954-1965.

Campos, P. F., & Gilbert, T. M. P. (2012). DNA extraction from formalin-fixed material. Ancient DNA, Humana Press, 81–85.

Clarke, F. E. (1899). Notes on New Zealand Galaxidae, more especially those of the western slopes: with descriptions of new species. Transactions and Proceedings of the New Zealand Institute, 31, 78–91.

Cooper, R. A., & Millener, P. R. (1993). The New Zealand biota: Historical background and new research. Trends in Ecology and Evolution, 8(12), 429–433.

Dabney, J., Knapp, M., Glocke, I., Gansauge, M. T., Weihmann, A., Nickel, B., …, & Meyer, M. (2013). Complete mitochondrial genome sequence of a Middle Pleistocene cave bear reconstructed from ultrashort DNA fragments. Proceedings of the National Academy of Sciences of the United States of America, 110(39), 15758–15763.

Darriba, D., Taboada, G. L., Doallo, R., & Posada, D. (2012). jModelTest 2: more models, new heuristics and parallel computing. Nature Methods, 9(8), 772–772.

Drummond, A. J., Suchard, M. A., Xie D., & Rambaut, A. (2012) Bayesian Phylogenetics with BEAUti and the BEAST 1.7. Molecular Biology and Evolution, 29(8), 1969–1973.

Easton, L. J., Tennyson, A. J., & Rawlence, N. J. (2021). A new species of Leiopelma frog (Amphibia: Anura: Leiopelmatidae) from the late Pliocene of New Zealand. New Zealand Journal of Zoology, 1–10.

Edgar, R. C. (2004). MUSCLE: A multiple sequence alignment method with reduced time and space complexity. BMC Bioinformatics, 5(1), 1–19.

Ferreira, M. A. R., & Suchard, M. A. (2008) Bayesian analysis of elapsed times in continuous-time Markov chains. Canadian Journal of Statistics, 36, 355–368.

Friedman, M., & DeSalle, R. (2008). Mitochondrial DNA extraction and sequencing of formalin-fixed archival snake tissue. DNA Sequence, 19(5), 433–437.

Fyfe, R., & Bradshaw, J. (2020). A review of the role of diadromous ikawai (freshwater fish) in the Māori economy and culture of Te Wai Pounamu (South Island), Aotearoa New Zealand. Records of the Canterbury Museum, 34, 35–55.

Gansauge, M. T., Gerber, T., Glocke, I., Korlević, P., Lippik, L., Nagel, S., … & Meyer, M. (2017). Single-stranded DNA library preparation from highly degraded DNA using T4 DNA ligase. Nucleic acids research, 45(10), e79–e79.

Gansauge, M. T., & Meyer, M. (2013). Single-stranded DNA library preparation for the sequencing of ancient or damaged DNA. Nature Protocols, 8(4), 737–748.

Gansauge, M. T., Aximu-Petri, A., Nagel, S., & Meyer, M. (2020). Manual and automated preparation of single-stranded DNA libraries for the sequencing of DNA from ancient biological remains and other sources of highly degraded DNA. Nature Protocols, 15(8), 2279–2300.

Garrigos, Y. E., Hugueny, B., Koerner, K., Ibanez, C., Bonillo, C., Pruvost, P., … & Gaubert, P. (2013). Non-invasive ancient DNA protocol for fluid-preserved specimens and phylogenetic systematics of the genus Orestias (Teleostei: Cyprinodontidae). Zootaxa, 3640(3), 373–394.

Ginolhac, A., Rasmussen, M., Gilbert, M. T. P., Willerslev, E., & Orlando, L. (2011). mapDamage: testing for damage patterns in ancient DNA sequences. Bioinformatics, 27(15), 2153–2155.

González Fortes, G., & Paijmans, J. L. (2019). Whole-genome capture of ancient DNA using homemade baits. In Ancient DNA (pp. 93–105). Humana Press, New York, NY.

Hahn, E. E., Alexander, M. R., Grealy, A., Stiller, J., Gardiner, D. M., & Holleley, C. E. (2021). Unlocking inaccessible historical genomes preserved in formalin. Molecular Ecology Resources.

Hammer, M. P., Adams, M., Unmack, P. J., & Walker, K. F. (2007). A rethink on Retropinna: conservation implications of new taxa and significant genetic substructure in Australian smelts (Pisces: Retropinnidae). Marine and Freshwater Research, 58(4), 327–341.

Hiroa, T. R. (1926). The Maori craft of netting. Transactions and Proceedings of the New Zealand Institute, 56, 597–646

Hitchmough, R., Bull, L., & Cromarty, P. (2002). New Zealand threat classification system lists. Threatened Species Occasional Publication, 23.

Hughes, J. M., Schmidt, D. J., Macdonald, J. I., Huey, J. A., & Crook, D. A. (2014). Low interbasin connectivity in a facultatively diadromous fish: evidence from genetics and otolith chemistry. Molecular Ecology, 23(5), 1000–1013.

Hykin, S. M., Bi, K., & McGuire, J. A. (2015). Fixing formalin: a method to recover genomic-scale DNA sequence data from formalin-fixed museum specimens using high-throughput sequencing. PloS One, 10(10), e0141579.

Johnson, G. D., & Patterson, C. (1996). Relationships of lower euteleostean fishes. Interrelationships of fishes. Academic Press, London.

Johnston, R. M. (1882). General and critical observations on the fishes of Tasmania; with a classified catalogue of all the known species. In Papers & Proceedings of the Royal Society of Tasmania (pp. 51–170).

Jónsson, H., Ginolhac, A., Schubert, M., Johnson, P. L., & Orlando, L. (2013). mapDamage2. 0: fast approximate Bayesian estimates of ancient DNA damage parameters. Bioinformatics, 29(13), 1682–1684.

Kaminskas, S. (2021). Alien pathogens and parasites impacting native freshwater fish of southern Australia: a scientific and historical review. Australian Zoologist, 41(4), 696–730.

Lanfear, R., Calcott, B., Ho, S. Y., & Guindon, S. (2012). PartitionFinder: combined selection of partitioning schemes and substitution models for phylogenetic analyses. Molecular Biology and Evolution, 29(6), 1695–1701.

Lanfear, R., Frandsen, P. B., Wright, A. M., Senfeld, T., & Calcott, B. (2017). PartitionFinder 2: new methods for selecting partitioned models of evolution for molecular and morphological phylogenetic analyses. Molecular Biology and Evolution, 34(3), 772–773.

Leach, F. (2006). Fishing in pre-european New Zealand. Archaeofauna, (15), 19–276.

Lee, D. E., McDowall, R. M., & Lindqvist, J. K. (2007). Galaxias fossils from Miocene lake deposits, Otago, New Zealand: The earliest records of the Southern Hemisphere family Galaxiidae (Teleostei). Journal of the Royal Society of New Zealand, 37(3), 109–130.

Lee, F., & Perry, G. L. (2019). Assessing the role of off-take and source–sink dynamics in the extinction of the amphidromous New Zealand grayling (Prototroctes oxyrhynchus). Freshwater Biology, 64(10), 1747–1754.

Leigh, J. W., & Bryant, D. (2015). POPART: full-feature software for haplotype network construction. Methods in Ecology and Evolution, 6(9), 1110–1116.

Li, H., & Durbin, R. (2009). Fast and accurate short read alignment with Burrows– Wheeler transform. Bioinformatics, 25(14), 1754–1760.

Li, H., Handsaker, B., Wysoker, A., Fennell, T., Ruan, J., Homer, N., … & Durbin, R. (2009). The sequence alignment/map format and SAMtools. Bioinformatics, 25(16), 2078–2079.

Li, J., Xia, R., McDowall, R. M., López, J. A., Lei, G., & Fu, C. (2010). Phylogenetic position of the enigmatic Lepidogalaxias salamandroides with comment on the orders of lower euteleostean fishes. Molecular Phylogenetics and Evolution, 57(2), 932–936.

McDowall, R. M. (1969). Relationships of galaxioid fishes with a further discussion of salmoniform classification. Copeia, 796–824.

McDowall, R. M. (1976). Fishes of the family Prototroctidae (Salmoniformes). Marine and Freshwater Research, 27(4), 641–659.

McDowall, R. M. (1990). When galaxiid and salmonid fishes meet–a family reunion in New Zealand. Journal of Fish Biology, 37, 35–43.

McDowall, R. M., Kennedy, E. M., & Alloway, B. V. (2006). A fossil southern grayling, genus Prototroctes, from the Pleistocene of north-eastern New Zealand (Teleostei: Retropinnidae). Journal of the Royal Society of New Zealand, 36(1), 27–36.

McDowall, R. M. (2011). Ikawai: freshwater fishes in Māori culture and economy. Canterbury University Press, Christchurch.

McKenna, A., Hanna, M., Banks, E., Sivachenko, A., Cibulskis, K., Kernytsky, A., … & DePristo, M. A. (2010). The Genome Analysis Toolkit: a MapReduce framework for analyzing next-generation DNA sequencing data. Genome Research, 20(9), 1297–1303.

McWethy, D. B., Wilmshurst, J. M., Whitlock, C., Wood, J. R., & McGlone, M. S. (2014). A high-resolution chronology of rapid forest transitions following Polynesian arrival in New Zealand. PLoS One, 9(11), e111328.

Mildenhall, D. C., Mortimer, N., Bassett, K. N., & Kennedy, E. M. (2014). Oligocene paleogeography of New Zealand: maximum marine transgression. New Zealand Journal of Geology and Geophysics, 57(2), 107–109.

Miya, M., & Nishida, M. (2000). Use of mitogenomic information in teleostean molecular phylogenetics: a tree-based exploration under the maximum-parsimony optimality criterion. Molecular Phylogenetics and Evolution, 17(3), 437–455.

Nelson, J. S. (1994). Fishes of the world. Wiley, New York.

Page, T. J., & Hughes, J. M. (2010). Comparing the performance of multiple mitochondrial genes in the analysis of Australian freshwater fishes. Journal of Fish Biology, 77(9), 2093–2122.

Phillipps, W. J. (1923). Life history of the New Zealand grayling, Prototroctes oxyrhynchus. The New Zealand Journal of Science and Technology, 6(11), 5–17.

Pierson, T. W., Kieran, T. J., Clause, A. G., & Castleberry, N. L. (2020). Preservation-induced morphological change in salamanders and failed DNA extraction from a decades-old museum specimen: implications for Plethodon ainsworthi. Journal of Herpetology, 54(2), 137–143.

Preston, C. (1986). From the archives. West Coast Acclimatisation Society Annual Report 1986, 16.

Pyron, R. A., Beamer, D. A., Holzheuser, C. R., Lemmon, E. M., Lemmon, A. R., Wynn, H., & O’Connell, K. A. (2022). Contextualizing enigmatic extinctions using genomic DNA from fluid-preserved museum specimens of Desmognathus salamanders. Conservation Genetics, 1–12.

Rambaut, A., Drummond, A. J., Xie, D., Baele, G., & Suchard, M. A. (2018). Posterior summarization in Bayesian phylogenetics using Tracer 1.7. Systematic Biology, 67(5), 901–904.

Rawlence, N. J., Collins, C. J., Anderson, C. N., Maxwell, J. J., Smith, I. W., Robertson, C., … & Waters, J. M. (2016). Human-mediated extirpation of the unique Chatham Islands sea lion and implications for the conservation management of remaining New Zealand sea lion populations. Molecular Ecology, 25(16), 3950–3961.

Rawlence, N. J., Verry, A., Greig, K., Maxwell, J. J., Shepherd, L. S. & Walter, R. (2020). Reconstructing the impact of humans on Aotearoa New Zealand’s biodiversity. In: Decocq, G. (ed) Historical Ecology. Springer, In Press.

Rozas, J., Ferrer-Mata, A., Sánchez-DelBarrio, J. C., Guirao-Rico, S., Librado, P., Ramos-Onsins, S. E., & Sánchez-Gracia, A. (2017). DnaSP 6: DNA sequence polymorphism analysis of large data sets. Molecular Biology and Evolution, 34(12), 3299–3302.

Rutland, J. (1877). On the habits of the New Zealand grayling. Transactions and Proceedings of the New Zealand Institute Art, XXXII, 250–252.

Saville-Kent, W. (1887). Observations on the acclimatisation of the true salmon (Salmo salar), in Tasmanian waters, and upon the reported salmon disease at the breeding establishment on the River Plenty. In Papers & Proceedings of the Royal Society of Tasmania (pp. 54–66).

Scarsbrook, L., Verry, A. J., Walton, K., Hitchmough, R. A., & Rawlence, N. J. (2022). Ancient mitochondrial genomes recovered from small vertebrate bones through minimally destructive DNA extraction: phylogeography of the New Zealand gecko genus Hoplodactylus. Molecular Ecology, 00, 1–0. https://doi.org/10.1111/mec.16434

Schmidt, D. J., Islam, M. R. U., & Hughes, J. M. (2016). Complete mitogenomes for two lineages of the Australian smelt, Retropinna semoni (Osmeriformes: Retropinnidae). Mitochondrial DNA Part B, 1(1), 615–616.

Schubert, M., Ermini, L., Sarkissian, C. D., Jónsson, H., Ginolhac, A., Schaefer, R., … & Orlando, L. (2014). Characterization of ancient and modern genomes by SNP detection and phylogenomic and metagenomic analysis using PALEOMIX. Nature Protocols, 9(5), 1056–1082.

Schubert, M., Lindgreen, S., & Orlando, L. (2016). AdapterRemoval v2: rapid adapter trimming, identification, and read merging. BMC Research Notes, 9(1), 1–7.

Schwarzhans, W., Scofield, R. P., Tennyson, A. J., Worthy, J. P., & Worthy, T. H. (2011). Fish remains, mostly otoliths, from the non-marine early Miocene of Otago, New Zealand. Acta Palaeontologica Polonica, 57(2), 319–350.

Schwarzhans, W. W., & Nielsen, S. N. (2021). Fish otoliths from the early Miocene of Chile: a window into the evolution of marine bony fishes in the Southeast Pacific. Swiss Journal of Palaeontology, 140(1), 1–62.

Seersholm, F. V., Cole, T. L., Grealy, A., Rawlence, N. J., Greig, K., Knapp, M., … & Bunce, M. (2018). Subsistence practices, past biodiversity, and anthropogenic impacts revealed by New Zealand-wide ancient DNA survey. Proceedings of the National Academy of Sciences, 115(30), 7771–7776.

Straube, N., Li, C., Mertzen, M., Yuan, H., & Moritz, T. (2018). A phylogenomic approach to reconstruct interrelationships of main clupeocephalan lineages with a critical discussion of morphological apomorphies. BMC Evolutionary Biology, 18(1), 1–17.

Thacker, C. E., Unmack, P. J., Matsui, L., & Rifenbark, N. (2007). Comparative phylogeography of five sympatric Hypseleotris species (Teleostei: Eleotridae) in southeastern Australia reveals a complex pattern of drainage basin exchanges with little congruence across species. Journal of Biogeography, 34(9), 1518–1533.

Waters, J. M., Saruwatari, T., Kobayashi, T., Oohara, I., McDowall, R. M., & Wallis, G. P. (2002). Phylogenetic placement of retropinnid fishes: data set incongruence can be reduced by using asymmetric character state transformation costs. Systematic Biology, 51(3), 432–449.

Waters, J. M., Fraser, C. I., & Hewitt, G. M. (2013). Founder takes all: density-dependent processes structure biodiversity. Trends in Ecology & Evolution, 28(2), 78–85.

Wilmshurst, J. M., Anderson, A. J., Higham, T. F., & Worthy, T. H. (2008). Dating the late prehistoric dispersal of Polynesians to New Zealand using the commensal Pacific rat. Proceedings of the National Academy of Sciences, 105(22), 7676–7680.

Wilson, M. V., & Williams, R. R. (1991). New Paleocene genus and species of smelt (Teleostei: Osmeridae) from freshwater deposits of the Paskapoo Formation, Alberta, Canada, and comments on osmerid phylogeny. Journal of Vertebrate Paleontology, 11(4), 434–451.

Worthy, T. H. (1987). Osteological observations on the larger species of the skink Cyclodina and the subfossil occurrence of these and the gecko Hoplodactylus duvaucelii in the North Island, New Zealand. New Zealand Journal of Zoology, 14(2), 219–229.

