## Supplementary Information for "Ancient DNA from the extinct New Zealand grayling (*Prototroctes oxyrhynchus*) reveals evidence for Miocene marine dispersal"

Prepared for submission to *Freshwater Biology*

### Table of Contents:



| | |
|---|---|
| **Supplementary Figures (S1-S2)** | Page 2 |
| **Supplementary Tables (S1-S3)** | Page 4 |

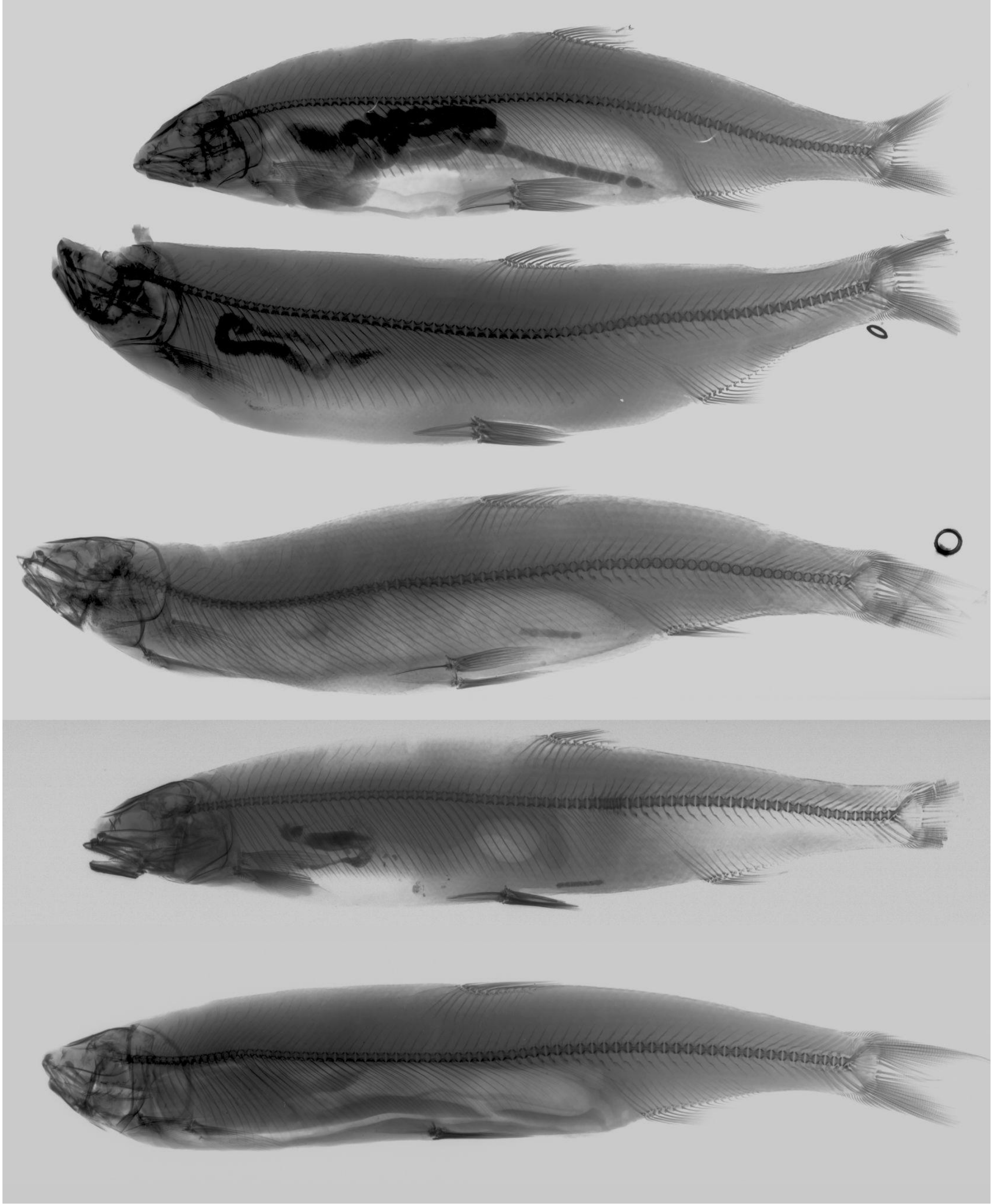

**Supplementary Figure 1** X-ray images of New Zealand grayling (*Prototroctes oxyrhynchus*) specimens at the British National History Museum (NMUK), used to assess preservation of calcareous otoliths. Images were generated by James Maclaine (NMUK). Top to bottom: 1870.5.22.17 (syntype); 1870.5.22.18 (syntype); 1873.12.13.69; 1886.11.18.80; 1935.3.14.65.

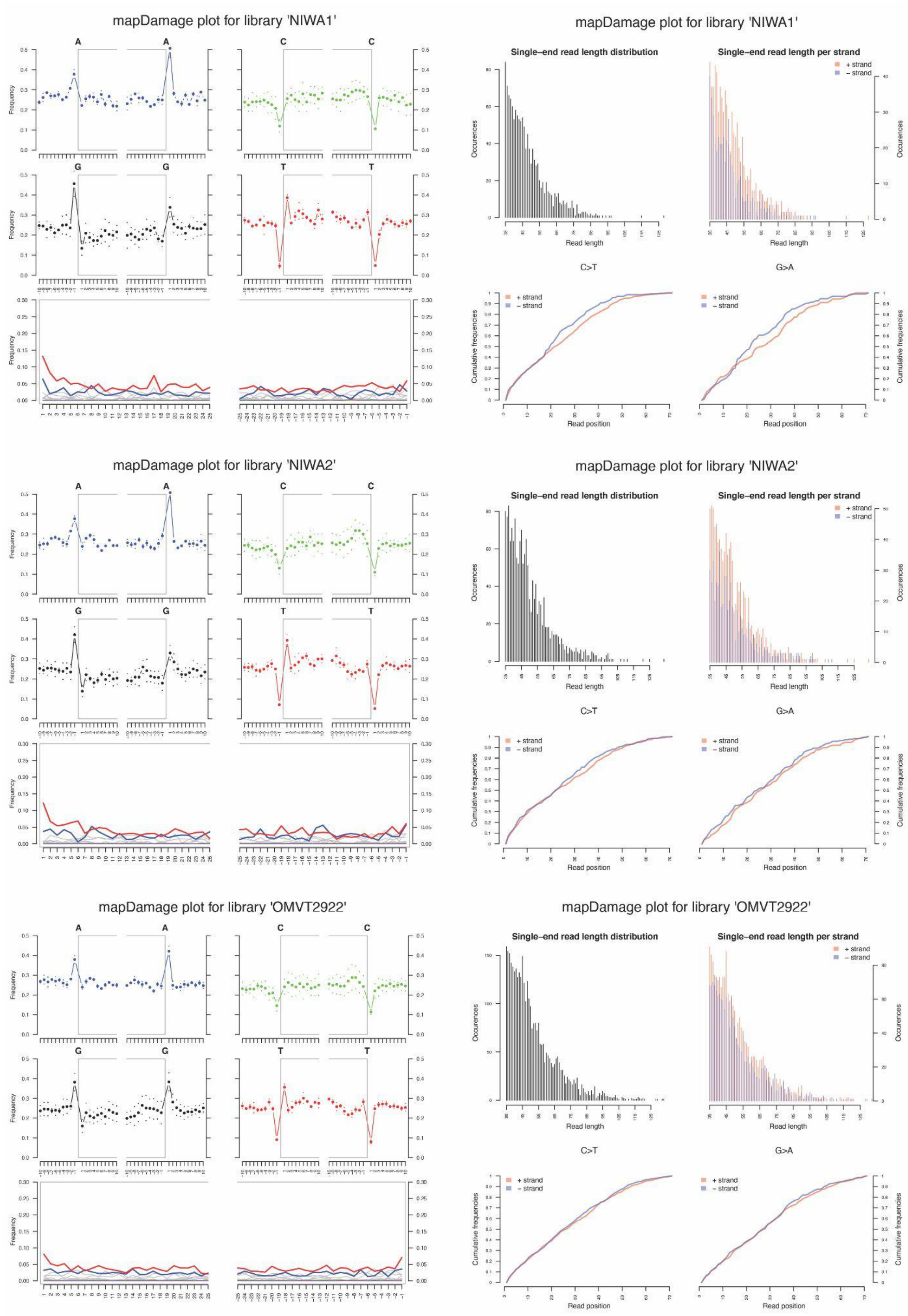


**Supplementary Figure 2** MapDamage reports for mapped collapsed reads of historic New Zealand grayling (*Prototroctes oxyrhynchus*) single-stranded libraries (NIWA1, NIWA2, OMVT2922). The left panels show characteristic high frequency of purines (A and G) at read termini (top) and accumulation of 5' C to T (red curve) misincorporations (bottom), which authenticate ancient DNA. The right panel shows characteristic short fragment length of single-end mapped reads (top).

**Supplementary Table 1** Sample information for New Zealand grayling (*Prototroctes oxyrhynchus*) specimens including: institution (Auckland War Memorial Museum (AWMM), Canterbury Museum (CM), Museum of New Zealand Te Papa Tongarewa (NMNZ), Otago Museum (OM) and unregistered (unreg.), sample identification (ID), tissue type and locality.

| Institution | Sample ID | Tissue Type | Locality |
|---|---|---|---|
| AWMM | MA110 | muscle ('wet') | New Zealand |
| AWMM | MA189 | muscle ('wet') | New Zealand |
| AWMM | MA190 | muscle ('wet') | New Zealand |
| AWMM | NGD01 | scale ('dried') | New Zealand |
| CM | CM1631 | muscle ('wet') | Westland |
| CM | CM1632 | muscle ('wet') | Waiau River, Southland |
| CM | CM1633 | muscle ('wet') | Orari River, Canterbury |
| NMNZ | NMNZ130 | scale ('dried') | Waimakariri River, Canterbury |
| NMNZ | NMNZP339 | muscle ('wet') | Turanganui River, Gisborne |
| OM | OMVT2922 | scale ('dried') | Clutha River, Otago |
| NIWA | NIWA1 | scale ('dried') | Hokitika River, Westland |
| NIWA | NIWA2 | scale ('dried') | Hokitika River, Westland |

**Supplementary Table 2** GenBank Accession numbers for Retropinnidae (R) and Osmeridae (O) sequences used in phylogenetic analyses. ***** denotes sequences derived through annotation extraction from complete mitochondrial genomes. Bolded accession numbers indicate sequences generated in this study.

| Species | Mitogenome | Cytochrome B | 16S rRNA |
|---|---|---|---|
| *Prototroctes maraena* (R) | **ON220597** | **ON161129–ON161135** | AF454844.1, HM151552.1 |
| *Prototroctes oxyrhynchus* (R) | **ON220594–ON220596** | - | - |
| *Retropinna retropinna* (R) | NC.004598.1 | *AP004108.1, *NC004598.1, FJ392549.1, FJ392550.1 | *AP004108.1, *NC004598.1, AF454842.1, FJ392551.1, FJ392552.1 |
| *Retropinna semoni* (R) | KX421784.1, KX421785.1 | *KX421784.1, *KX421785.1, NC031372.1, HM007065.1–HM007067.1, JN232588.1, JX914039.1–JX914083.1, MG867590.1–MG867657.1 | *KX421784.1, *KX421785.1, *NC031372.1, AF454845.1 |
| *Retropinna tasmanica* (R) | - | AF112321.1, JN232589.1 | AF112342.1 |
| *Stokellia anisodon* (R) | - | JN232590.1 | AF454843.1, HM151553.1 |
| *Hypomesus japonicus* (O) | MH636616.1 | - | - |
| *Hypomesus nipponensis* (O) | HM106489.1 | - | - |
| *Hypomesus olidus* (O) | KP281293.1 | - | - |
| *Mallotus villosus* (O) | HM106491.1 | - | - |
| *Osmerus eperlanus* (O) | NC.052758.1 | - | - |
| *Osmerus mordax* (O) | HM106493.1 | - | - |
| *Plecoglossus altivelis* (O) | AB047553.2 | - | - |
| *Salangichthys microdon* (O) | NC.004599.1 | - | - |
| *Salanx ariakensis* (O) | AP006231.1, KM517200.1 | - | - |
| *Salanx chinensis* (O) | MW131880.1 | - | - |
| *Salanx cuvieri* (O) | KJ645978.1 | - | - |

**Supplementary Table 3** Mitochondrial genome assembly statistics for historic New Zealand grayling (*Prototroctes oxyrhynchus*) specimens (see Supplementary Table 1) with reads generated through single-stranded library preparation. Summary statistics are not reported (denoted by "-") for "GC content (%)", "Ambiguous Sites", "Ambiguous bases (%)", "Reference sequence coverage (%)" and "Contig length (bp)" in the majority of individuals given insufficient reference sequence coverage.

| Sample | Paired-end reads | Collapsed reads | Mapped reads (raw) | Mapped reads (excl. duplicates) | Endogenous DNA (%) | Mean fragment length (bp) | GC content (%) | Mean coverage ± SD | Ambiguous Sites | Ambiguous bases (%) | Reference sequence coverage (%) | Contig length (bp) |
|---|---|---|---|---|---|---|---|---|---|---|---|---|
| NGD01 | 4,671,869 | 2,020,413 | 58 | 58 | 0.00% | 43.7 | - | 0.15 | - | - | - | - |
| NIWA1 | 5,080,301 | 1,686,400 | 1,310 | 1,262 | 0.10% | 47.7 | 47.70% | 3.63 | 6,418 | 38.68% | 79.80% | 16,593 |
| NIWA2 | 5,412,541 | 2,046,902 | 1,652 | 1,582 | 0.10% | 49.7 | 47.60% | 4.74 | 4,636 | 27.94% | 86.60% | 16,593 |
| NMNZ130 | 4,584,116 | 1,935,302 | 125 | 115 | 0.00% | 43.5 | - | 0.3 | - | - | - | - |
| OMVT2922 | 5,514,150 | 2,015,039 | 3,659 | 3,501 | 0.20% | 51.4 | 48.10% | 10.84 | 711 | 4.29% | 96.20% | 16,592 |
| CM1631 | 6,545,738 | 2,110,943 | 11 | 1 | 0.00% | 38 | - | 0 | - | - | - | - |
| CM1632 | 8,947,473 | 3,908,601 | 0 | 0 | 0.00% | 0 | - | 0 | - | - | - | - |
| CM1633 | 7,885,774 | 3,263,917 | 0 | 0 | 0.00% | 0 | - | 0 | - | - | - | - |
| MA110 | 5,808,531 | 1,841,448 | 10 | 7 | 0.00% | 49.9 | - | 0.02 | - | - | - | - |
| MA189 | 6,157,394 | 2,588,468 | 0 | 0 | 0.00% | 0 | - | 0 | - | - | - | - |
| MA190 | 4,805,344 | 1,843,397 | 24 | 2 | 0.00% | 57.5 | - | 0.01 | - | - | - | - |
| NMNZP339 | 4,798,497 | 1,642,070 | 0 | 0 | 0.00% | 0 | - | 0 | - | - | - | - |
